# Learning from the physical consequences of our actions improves motor memory

**DOI:** 10.1101/2021.10.07.463541

**Authors:** Amanda Bakkum, Daniel S. Marigold

## Abstract

Actions have consequences. Motor learning involves correcting actions that lead to movement errors and remembering these actions for future behavior. In most laboratory situations, movement errors have no physical consequences and simply indicate the progress of learning. Here we asked how experiencing a physical consequence when making a movement error affects motor learning. Two groups of participants adapted to a new, prism-induced mapping between visual input and motor output while performing a precision walking task. Importantly, one group experienced an unexpected slip perturbation when making foot-placement errors during adaptation. Because of our innate drive for safety, and the fact that balance is fundamental to movement, we hypothesized that this experience would enhance motor memory. Learning generalized to different walking tasks to a greater extent in the group who experienced the adverse physical consequence. This group also showed faster relearning one week later despite exposure to a competing mapping during initial learning—evidence of greater memory consolidation. The group differences in generalization and consolidation occurred even though they both experienced similar magnitude foot-placement errors and adapted at similar rates. Our results suggest the brain considers the potential physical consequences of movement error when learning and that balance-threatening consequences serve to enhance this process.

## INTRODUCTION

Every action has a consequence. Different factors, such as reward and punishment, can serve to strengthen or reinforce the association between actions and their consequences and are therefore compelling modulators of behavior (Abe et al. 2011; Galea et al. 2015; Thorndike 1933; Skinner 1938). Motor learning involves correcting actions that lead to errors and remembering these actions for future performance. Sensory feedback plays an important role in this ability (Henriques and Cressman 2012; Maeda et al. 2017a), though research also shows that punishing errors can accelerate motor learning whereas rewarding movement accuracy is beneficial for retaining motor memories (Abe et al. 2011; Galea et al. 2015; Hill et al. 2020; Quattrocchi et al. 2018; Song and Smiley-Oyen 2017; Wächter et al. 2009). These experiments used monetary reward and punishment to reinforce learning, which does not reflect the movement consequences we experience in daily life. Rather, errors in everyday goal-directed movement often lead to physical consequences.

Some physical consequences of movement error are benign, while others have the potential to cause harm to the individual. An accidental misstep off a sidewalk, for example, can lead to an injurious fall. Thus, movement decisions are often made with personal safety in mind. Given that many daily movements are plagued by inherent stability challenges, a primary concern of the nervous system is to maintain balance. Even moderate perceived threats to postural stability elicit movement strategies that serve to safeguard balance (Adkin et al. 2000; Adkin and Carpenter 2018; Brown et al. 2002; Manista and Ahmed 2012). Such safety-driven movement strategies are also observed when walking across unstable terrain (Marigold and Patla 2002). How might experiencing a balance-threatening physical consequence when making movement errors affect motor learning?

Evidence suggests that experiencing an unpleasant or dangerous physical consequence, or the threat of these types of consequences, can influence learning and memory. For example, rodents can quickly learn and subsequently remember the spatial location where foot shocks occur in an environment (Stuchlík et al. 2013; Willis et al. 2017). In addition, in humans, pairing an electric shock with images of neutral objects improves item recognition memory (Dunsmoor et al. 2015; Starita et al. 2019). Even the threat of being shocked enhances declarative memory (Murty et al. 2012). Because of our innate drive for safety, and the fact that balance is fundamental to movement, we hypothesized that experiencing a balance-threatening physical consequence when making a movement error would enhance motor memory.

To test this hypothesis, we had two groups of participants adapt to a new visuomotor mapping induced by prism lenses while performing a precision walking task that required them to step accurately to a target (Bakkum et al. 2020, 2021; Maeda et al. 2017a). Prism lenses cause errors in movement because they alter the normal relationship (or mapping) between visual input and motor output. Learning this mapping is thus essential for achieving movement accuracy. The groups differed in terms of the consequence experienced when making foot-placement errors to the target. Specifically, one group experienced a balance-threatening slip perturbation caused by stepping on a concealed slippery surface positioned adjacent to the target. In contrast to an artificial consequence, like monetary gains or losses, this slip perturbation is an adverse physical consequence that can occur in daily life.

We determined how the balance-threatening physical consequence affected 1) generalization of the learned visuomotor mapping across different visually guided walking tasks, and 2) consolidation of the learned mapping one week later. Generalization and consolidation are two hallmarks of motor memory (Kitago and Krakauer 2013, Krakauer et al. 2019; Poggio and Bizzi 2004). However, past sensorimotor adaptation research demonstrates mixed evidence of support for both generalization (Alexander et al. 2011, 2013; Ghahramani et al. 1996; Krakauer et al. 2000; Wang 2008) and motor memory consolidation (Brashers-Krug et al. 1996; Caithness et al. 2004; Krakauer et al. 2005; Maeda et al. 2017b, 2018; Malone et al. 2011). Here, we found that the group who experienced the adverse physical consequence better generalized learning to different walking tasks and showed greater consolidation. These results suggest our motor systems are tuned to remember motor behaviors that promote personal safety. These results may also help shape new neurorehabilitation strategies.

## MATERIALS AND METHODS

### Participants

Twenty-four participants (mean age ± SD = 26.4 ± 5.0 years; 11 men, 13 women) with no known musculoskeletal, visual (six participants wore corrective lenses or glasses), or neurological disease participated in this study. We randomly assigned these participants to one of two groups (n = 12 each; detailed below). We did not perform any a priori power analysis to determine sample size. Rather, we used sample sizes typical in the literature for this type of research (approximately 6 to 14 participants per group). The Office of Research Ethics at Simon Fraser University approved the study protocol, and all participants provided written informed consent prior to their participation.

### Experimental tasks and data collection

All participants adapted to a novel visuomotor mapping induced by prism lenses (Fig. 1a) while performing a precision walking task (Fig. 1b). For this task, participants stood at the beginning of the walkway (∼6 m in length) and waited for a go-cue to signal the start of each trial. Once cued, participants took a minimum of two steps before stepping with their right foot onto the medial-lateral (ML) center of a projected target (3 × 36 cm) without stopping. We instructed participants to be as accurate as possible in the ML dimension when stepping to the target. We used a long target to reduce the demand for accuracy in the anterior-posterior (AP) dimension and to prevent participants from using shuffle steps as they approached the target. We displayed the target in the center of the walking path using an LCD projector (Epson PowerLight 5535U; brightness of 5500 lumens). Participants performed the task under reduced light conditions (∼0.9 lux) to minimize the influence of environmental references and increase the visibility of the projected target.

**Figure 1.**
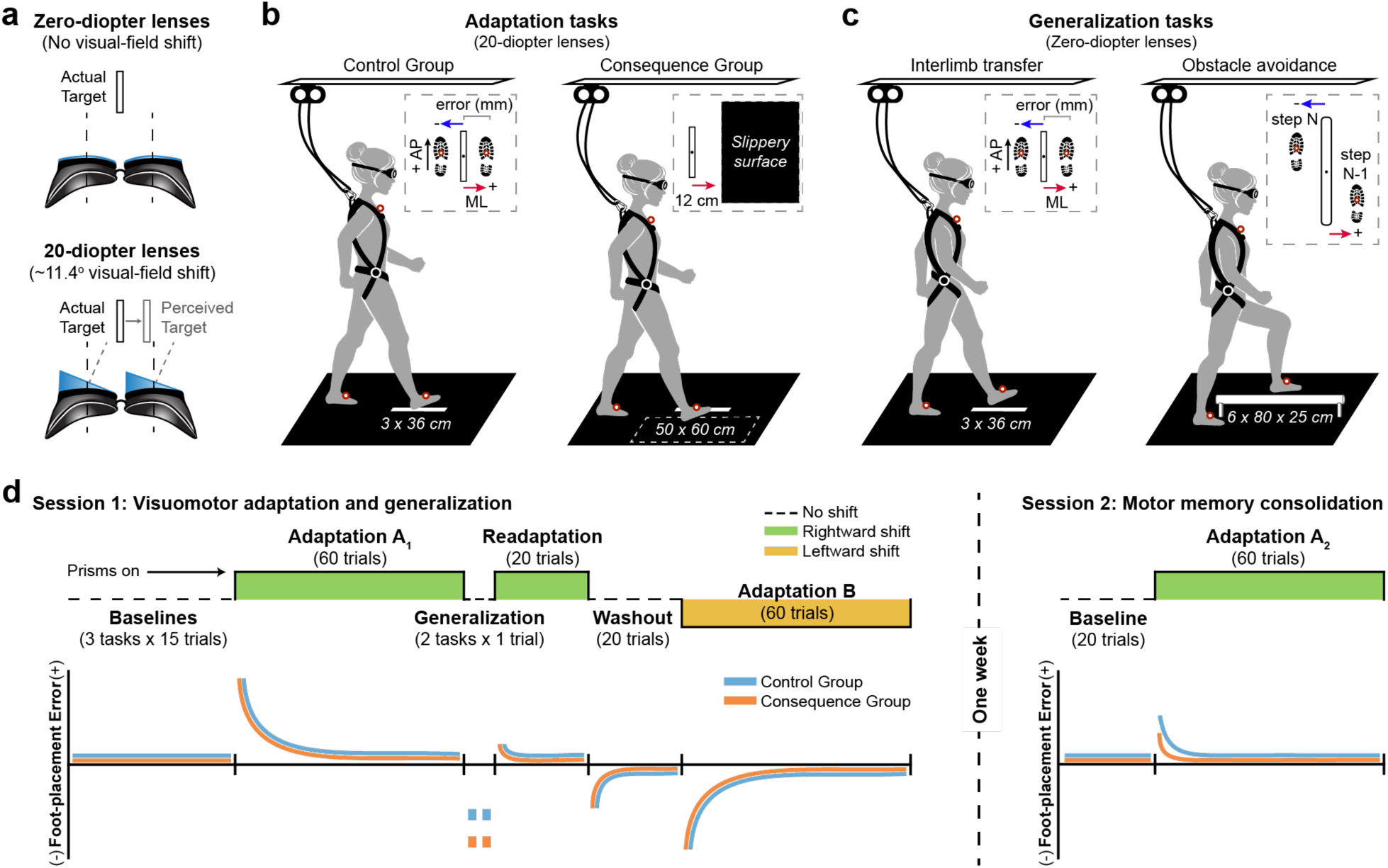
Experimental tasks and protocol. ***a)*** A simulated view of the stepping target through the goggles coupled with zero-diopter (non-visual-field-shifting) lenses and 20-diopter prism lenses that shift the perceived location of the target 11.4° to the right. ***b)*** An illustration of the precision walking task without and with an adverse physical consequence—a slippery surface—present next to the stepping target. Inset (left): a diagram showing positive (+) and negative (-) medial-lateral (ML) foot-placement error, defined as the distance between the position marker on the mid-foot and the center of the target. AP, anterior-posterior dimension in laboratory space. Inset (right): bird’s eye view of the location of the slippery surface relative to the target. ***c)*** An illustration of the generalization tasks. Left: interlimb transfer test. Note that the left foot is used to step to the target in this task. Inset: identical to that shown in (c). Right: obstacle avoidance task. Inset: a diagram showing positive (+) and negative (-) ML deviation from the obstacle, defined as the distance between the center of the obstacle and the position marker on the mid-foot of both the trailing foot (i.e., step N-1: right foot) and leading foot (i.e., step N: left foot). ***d)*** An illustration of the experimental protocol across both testing sessions (top), as well as predicted foot-placement error profiles for each phase of testing (bottom). During the first testing session, all participants performed baseline, adaptation, generalization, readaptation, and washout phases. The baseline phase included the precision walking task present in the adaptation phase and the two generalization walking tasks. Depending on the phase, participants wore goggles paired with either zero-diopter or 20-diopter lenses. To assess consolidation, participants repeated baseline and adaptation phases one week later. See text for details.

We tested 1) generalization of the learned visuomotor mapping across different visually guided walking tasks, and 2) consolidation of the learned mapping one week later. We assessed generalization during an interlimb transfer test and obstacle-avoidance task performed without the prism lenses. Interlimb transfer tests are commonly used in reaching experiments. The ability to negotiate over or around obstacles (e.g., stairs, curbs, toys on the ground) is essential for safe mobility. We have also previously found generalization to an obstacle-avoidance task (Alexander et al. 2011, 2013). During the interlimb transfer test, participants performed a single trial of the precision walking task using their left foot instead of their right foot to step to the target (Fig. 1c). For the obstacle avoidance task, participants walked along the same 6 m long path towards the right side of an obstacle (width = 6 cm; length = 80 cm; height = 25 cm) positioned in the center of the walkway. Once participants were beside the obstacle, they stepped laterally over the middle of it, first with their left leg (i.e., the leading leg), then their right leg (i.e., the trailing leg), before continuing to walk for several more steps (Fig. 1c). We instructed participants to avoid colliding with the obstacle.

We tracked body motion from infrared-emitting position markers placed on the participant’s chest (in line with the sternum) and bilaterally on the mid-foot (second to third metatarsal head) at 120 Hz during each task using an Optotrak Certus motion capture camera (Northern Digital, Waterloo, ON, Canada). To mitigate adaptation between trials, we instructed participants to only have their eyes open when they performed the walking tasks. An experimenter guided the participants back to the beginning of the walkway between trials while their eyes were closed. To prevent participants from learning a specific stepping sequence and increase the demand for visual feedback, we randomized the AP starting location (between 1.5 - 2.5 m) for each trial. We also encouraged participants to perform each task at a quick and constant pace to minimize online corrections of leg trajectory during the step to the target. Participants walked with an average speed (±SD) of 1.56 ± 0.16 m/s, and we later verified the absence of these corrections by looking at the kinematic profiles of the midfoot position markers. An experimenter demonstrated all tasks at the beginning of each testing session. Participants wore a safety harness at all times to prevent falling to the ground. No participant engaged the system during the experiment.

### Experimental protocol

We measured sensorimotor adaptation, generalization, and consolidation over two testing sessions, separated by one week. Figure 1d illustrates the experimental protocol and the predicted foot-placement error responses for each phase of testing. Depending on the phase, participants wore goggles coupled with either zero-diopter (non-visual-field-shifting) or 20-diopter prism lenses (Fig. 1a). The goggles block part of the peripheral visual field and participants had no option but to look through the lenses during each task. Participants performed three baseline phases (15 trials each), one for each visually guided walking task, while wearing the zero-diopter (i.e., non-visual-field-shifting) lenses. Participants performed the baseline phase for the adaptation task last, just prior to the adaptation phase. We counterbalanced the order of the two remaining baseline tasks for each participant and matched this order for the generalization tasks.

During the first adaptation phase (Adaptation A_1_), participants learned a new visuomotor mapping induced by the 20-diopter prism lenses (Fig. 1a) while performing 60 trials of the precision walking task using their right foot to step to the target. Participants adapted to the new visuomotor mapping with (consequence group; n = 12) or without (control group; n = 12) the possibility of experiencing an unexpected slip perturbation when making foot-placement errors. For the consequence group, we positioned a low-friction, polypropylene surface (50 × 60 cm) to the right of the target during prism exposure (Fig. 1b). We concealed this slippery surface using a solid black, low-friction fabric that covered the entire walking path. Exposure to the prism lenses induced a rightward deviation in foot placement to the target. This increased the likelihood of participants in the consequence group making contact with the slippery surface. On contact, the shear forces under the participants’ shoe at heel strike cause the low-friction fabric to slide over the slippery surface (kinetic coefficient of friction ≈ 0.09 μ_k_; for reference: ice ≈ 0.02 μ_k_) and require a reactive response to prevent falling. Participants only experienced the slip perturbation during stepping errors that were large enough that the foot made contact with the slippery surface. To prevent participants from being penalized during stepping errors within the normal range of late prism adaptation, we positioned the slippery surface 12 cm from the center of the target (Fig. 1b). A textured polyvinyl chloride bottom prevented the slippery surface from sliding along the walkway during foot contact.

Following adaptation to mapping A, participants performed the interlimb transfer test and the obstacle avoidance task (Fig. 1c,d) without the prism lenses to determine if the learned mapping was applied to the non-adapted tasks. Participants then performed 20 readaptation trials while wearing the rightward-shifting 20-diopter lenses to mitigate any deadaptation that might have occurred during the generalization phase. To confirm whether the learned mapping was stored, participants performed 20 washout trials of the adaptation task with the zero-diopter lenses. Finally, ∼15 minutes after adaptation, participants performed 60 trials of the same adaptation task (i.e., precision walking with the right foot stepping on the target) while wearing 20-diopter lenses that shifted the visual field in the opposite (i.e., leftward) direction of mapping A—we refer to this as mapping B (Fig. 1d). Following the initial testing session, the participants returned to the lab one week later so we could probe motor memory consolidation. We define consolidation as memory stabilization of the learned prism-induced mapping such that it is resistant to retrograde interference by another (competing) mapping (Krakauer et al. 2005). Participants first performed 20 baseline trials of the adaptation task while wearing the zero-diopter lenses. Thereafter, all participants repeated the 60 adaptation trials with the 20-diopter prism lenses to assess consolidation of mapping A. There was no slippery surface present for either group during the second testing session.

### Data and statistical analyses

We analyzed kinematic data (filtered using a fourth-order, low-pass Butterworth algorithm with a cut-off frequency of 6 Hz) using MATLAB (The MathWorks, Natick, MA) to calculate foot placement during the precision walking and obstacle avoidance tasks. We determined foot placement during each task as the moment of heel strike, derived using the vertical velocity of the mid-foot markers (Bakkum et al. 2020; O’Connor et al. 2007). For the precision walking task, we defined foot-placement error during the step to the target as the ML distance between the position of the mid-foot marker at heel strike and the center of the target. A positive value represents errors in the direction of the prism shift (i.e., to the right of the target) and negative values represent errors in the opposite direction to the prism shift (Fig. 1b). For the obstacle avoidance task, we calculated the ML distance between the obstacle and both the trailing foot (i.e., step N-1: right foot) and leading foot (i.e., step N: left foot) at heel strike using the mid-foot marker on each foot (Fig. 1c). For step N-1, increasing positive values represent a greater deviation of the right foot from the obstacle, whereas for step N, increasing negative values indicate greater deviation of the left foot from the obstacle.

During slip perturbations, we expected to see greater forward displacement and velocity of the right foot compared to baseline. Therefore, to test whether our hidden surface was effective at eliciting a slip, we calculated two measures for baseline and adaptation phase trials to quantify slip severity: slip distance and peak slip velocity. We calculated slip distance during the step to the target as the total AP displacement travelled by the right mid-foot marker between heel strike and slip end, the latter of which we defined as the moment AP velocity of the right mid-foot marker profile stabilized to zero. Note that AP displacement and velocity of the mid-foot marker has not stabilized to zero at heel strike; thus, we see a non-zero slip distance/velocity even for non-slip trials. We then calculated peak slip velocity as the maximum AP velocity of the right mid-foot marker within that same time interval (i.e., heel strike to slip end). We defined a slip perturbation trial, for each participant individually, as a slip distance or peak slip velocity greater than the mean plus two standard deviations of the last ten baseline trials. To determine differences in slip severity, we compared slip distance and slip velocity during the baseline phase (mean of the last ten trials) and the first adaptation trial between groups using separate two-way (Group x Phase) mixed-model ANOVAs, where we included participant as a random effect.

To determine how the adverse physical consequence associated with making an error affected adaptation, we compared foot-placement error during the baseline phase (mean of the last ten trials), first adaptation trial, late adaptation (mean of the last ten trials), and the first washout trial in the first testing session using a two-way (Group x Phase) mixed-model ANOVA (with participant as a random effect). When checking for the assumptions of an ANOVA, we found a potential outlier for the control group (studentized residual = 4.5). Excluding this data point did not change the results, suggesting it was non-influential. Thus, we included this data point in the final statistical model.

To determine if the learned visuomotor mapping generalized to the non-adapted limb during the precision walking task, we compared foot-placement error during the baseline phase when using the left foot to step to the target (mean of the last ten trials) and the generalization trial with a two-way (Group x Phase) mixed-model ANOVA (with participant as a random effect). Foot-placement errors in the direction opposite to the learned prism shift (i.e., a negative value) indicate generalization during the precision walking task. To determine if the learned visuomotor mapping generalized to the obstacle avoidance task, we compared foot-placement deviation from the obstacle during the baseline phase of this task (mean of the last ten trials) and the generalization trial for both the trailing foot (i.e., step N-1: right foot) and leading foot (i.e., step N: left foot). Foot placement relative to the obstacle in the direction opposite to the learned prism shift (i.e., to the left) indicates generalization during the obstacle avoidance task. Thus, for step N-1 (right foot), a smaller value reflects less deviation of the foot from the obstacle (i.e., a leftward shift in foot placement; Fig. 1c) and indicates generalization of the learned mapping to the trailing foot. For step N (left foot), a greater negative value reflects greater deviation of the foot from the obstacle (i.e., a leftward shift in foot-placement; Fig. 1c) and indicates generalization of the learned mapping to the leading foot. We used separate two-way (Group x Phase) mixed-model ANOVAs (with participant as a random effect) to determine if the learned mapping generalized to the obstacle avoidance task.

To determine the presence of consolidation, we quantified three measures: the first adaptation trial error, early adaptation error (i.e., mean of adaptation trials 2 – 8), and rate of adaptation. The first adaptation trial error represents the initial recall of the mapping. Early adaptation error captures the rapid reduction in foot-placement error early in the adaptation phase and does not assume that participants follow a specific pattern (i.e., it is a model-free measure) (Maeda et al. 2017b; Malone et al. 2011; Roemmich and Bastian 2015). A faster reduction in foot-placement error (i.e., faster relearning of the mapping, or savings) indicates consolidation of the learned mapping. Since model-based measures are also commonly used to quantify adaptation and savings (Criscimagna-Hemminger and Shadmehr 2008; Morton and Bastian 2004), we also calculated the rate of adaptation. This involved fitting an exponential model to the foot-placement error data during the 60 adaptation trials associated with mapping A for each testing session using the following equation:

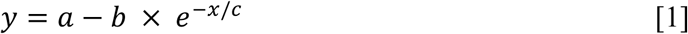

where, *a* is the residual error after asymptote (i.e., steady state), *b* is the magnitude of the adaptation required to reach *a* from the first trial, *c* is the decay constant, and *x* is the trial number. We defined the rate of adaptation as the number of trials taken to reach ∼63.2% of adaptation (Martin et al. 1996). We used separate two-way (Group x Session) mixed-model ANOVAs (with participant as a random effect) to determine differences in first adaptation trial error, early adaptation error, and adaptation rates between groups.

We used JMP 15 software (SAS Institute Inc., Cary, NC) with an alpha level of 0.05 for all statistical analyses. For ANOVAs, we used Tukey’s post hoc tests, where appropriate, when we found significant main effects or interactions. We report effect sizes as 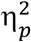. Post hoc test details (mean differences, 95% confidence intervals, and p values) are shown in supplementary material.

## RESULTS

### Contact with the slippery surface elicited a slip perturbation

To confirm that participants in the consequence group experienced a slip perturbation when missing the target, we calculated measures of slip distance and slip velocity during the baseline and adaptation phases for the consequence and control groups. Figure 2 illustrates group mean slip distance and peak slip velocity. We found that every participant in the consequence group (n = 12) experienced a slip during the first adaptation trial, which we define, for each participant individually, as a slip distance or peak slip velocity greater than the mean plus two standard deviations of the last ten trials of the baseline phase. During the first adaptation trial, the consequence group demonstrated a significantly greater slip distance compared to baseline and the control group (Fig. 2a and Fig. S1; mixed-model ANOVA, Group x Phase interaction: F_1,22_ = 85.49, p = 4.927e-9,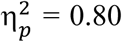), reflecting greater forward displacement of the right foot after contact with the slippery surface. Similarly, we found that peak slip velocity was significantly greater for the consequence group during the first adaptation trial compared to their baseline trials and the control group (Fig. 2b and Fig. S1; mixed-model ANOVA, Group x Phase interaction: F_1,22_ = 34.85, p = 6.103e-6, 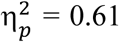). Additionally, all participants in the consequence group slipped during the second adaptation trial. Thereafter, the number of slips declined, and no participants slipped after the sixth adaptation trial. We found that the number of slips per trial differed slightly depending on the slip measure (i.e., slip distance or slip velocity), though this is likely due to the variability of the peak slip velocity measure. Overall, contact with the slippery surface during the precision walking task successfully elicited an adverse physical consequence, that is, a slip perturbation.

**Figure 2.**
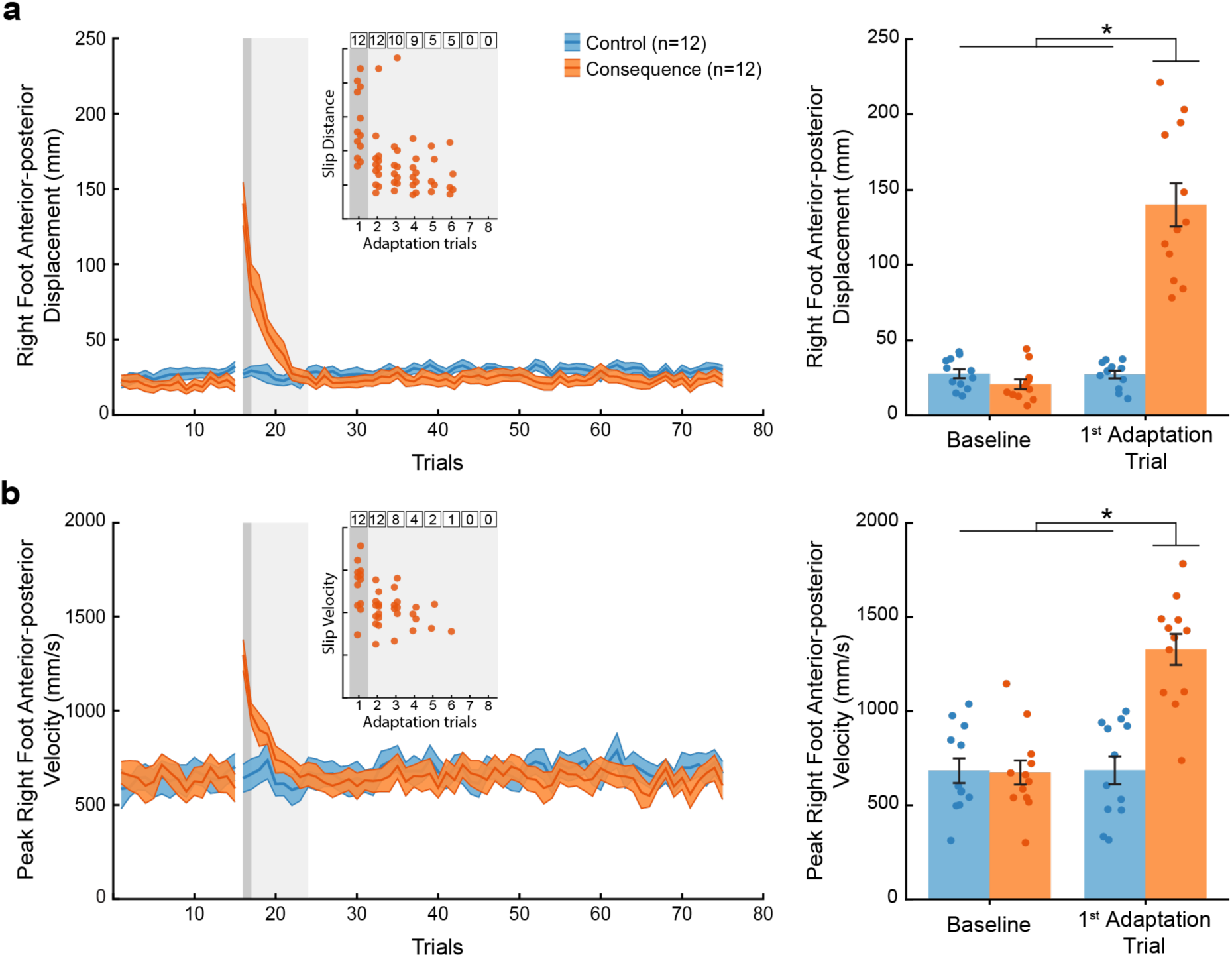
Slip measures. ***a)*** Left: Group mean ± SE slip distance across all trials for baseline and adaptation phases for the control (blue) and consequence (orange) groups. Right: Group mean ± SE slip distance for the baseline phase (mean of the last ten trials) and the first adaptation trial for the control (blue) and consequence (orange) groups. ***b)*** Left: Group mean ± SE peak slip velocity across all trials for baseline and adaptation phases for the control (blue) and consequence (orange) groups. Right: Group mean ± SE peak slip velocity for the baseline phase (mean of the last ten trials) and the first adaptation trial for the control (blue) and consequence (orange) groups. Individual participant values are superimposed. * Indicate that values are significantly different from each other based on post hoc tests (p < 0.05). See Fig. S1 for more detailed post hoc test results. Insets: individual data points showing significant evidence of a slip perturbation, defined as a slip distance or slip velocity of greater than the mean plus two standard deviations of the last ten baseline trials. The dark grey shaded box represents the first adaptation trial, and the light grey shaded box represents early adaptation trials. The numbered black boxes represent how many participants slipped during each trial. Every participant in the consequence group slipped during the first two adaptation trials. No participants slipped after the sixth adaptation trial.

### The presence of the adverse physical consequence did not disrupt initial visuomotor adaptation

Upon initial exposure to the prisms, all participants demonstrated a large, rightward deviation in foot placement relative to the target during the precision walking task. As participants adapted to the new, prism-induced visuomotor mapping, foot-placement error gradually returned to near-baseline levels of performance. Upon removal of the prism lenses, participants demonstrated a large, leftward deviation in foot-placement error (i.e., a negative aftereffect). These results are illustrated in Figure 3a.

**Figure 3.**
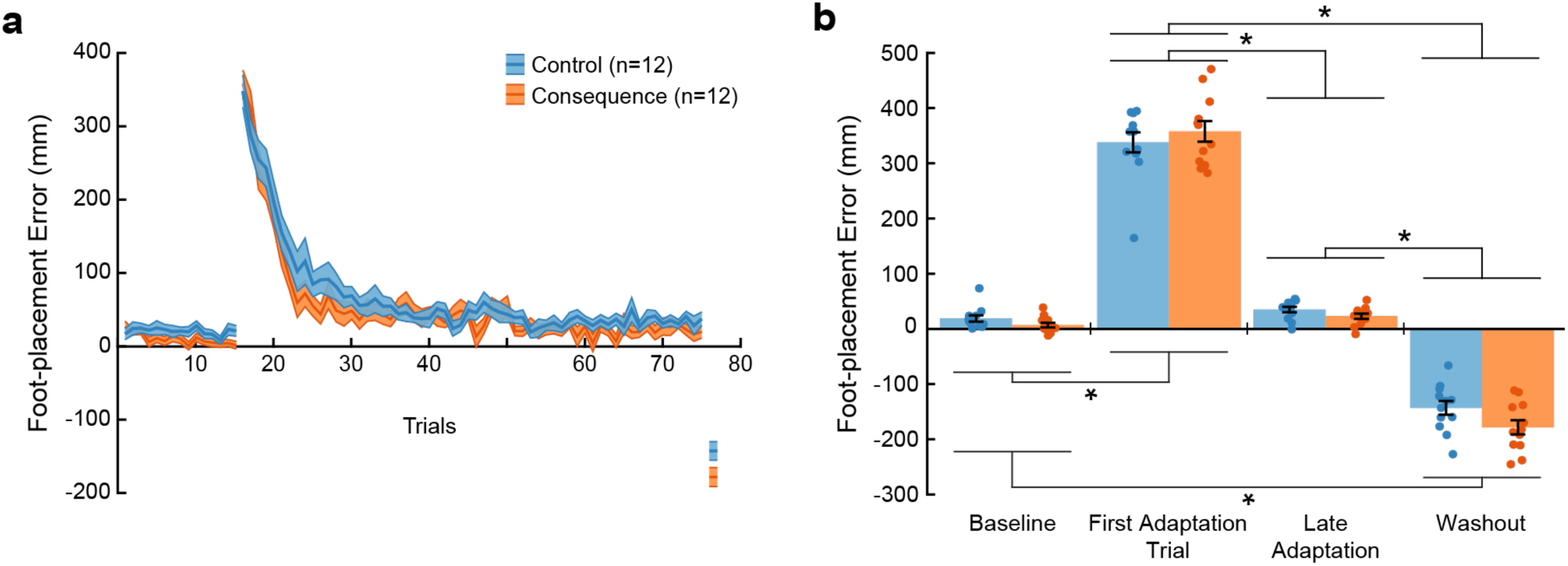
Visuomotor adaptation during session 1. ***a)*** Group mean ± SE foot-placement error across all trials for baseline and adaptation phases and the first washout trial during the first testing session for the control (blue) and consequence (orange) groups. ***b)*** Group mean ± SE foot-placement error for the baseline phase (mean of the last ten trials), first adaptation trial, late adaptation (mean of the last ten trials), and first washout trial during the first testing session for the control (blue) and consequence (orange) groups. Individual participant values are superimposed (n = 12 per group). A positive value represents errors in the direction of the prism shift (i.e., to the right of the target) and negative values represent errors in the opposite direction to the prism shift. * Indicate that values are significantly different from each other based on post hoc tests (p < 0.05). See Fig. S2 for more detailed post hoc test results.

To determine the effects of the adverse physical consequence on visuomotor adaptation, we compared foot-placement error across the baseline phase (mean of last ten trials), first adaptation trial, late adaptation (mean of last ten trials), and the first washout trial across groups during the first testing session. We found that foot-placement error differed depending on the phase (Fig. 3b; mixed-model ANOVA, Phase main effect: F_3,88_ = 657.4, p = 4.11e-60, 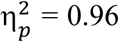). Post hoc tests indicated significantly greater foot-placement error during the first adaptation trial compared to the other phases (see also Fig. S2). Furthermore, foot-placement error during the first washout trial differed significantly from the other testing phases. We did not detect significant differences between the control and consequence groups across the testing phases. Overall, the adverse physical consequence experienced when making foot-placement errors did not affect the ability to adapt to the new, prism-induced visuomotor mapping.

### Learning generalized to a greater extent in the group who experienced the adverse physical consequence

To determine if the learned mapping generalized to non-adapted tasks, we had participants perform an interlimb transfer test and an obstacle-avoidance task following the initial adaptation to mapping A (Fig. 1c,d). During the interlimb transfer test, participants performed a single trial of the precision walking task using their left foot instead of their right foot to step to the target. For the obstacle avoidance task, participants walked toward the right side of an obstacle positioned in the center of the walkway and subsequently stepped laterally over it, first with their left leg (i.e., the leading leg), then their right leg (i.e., the trailing leg), before walking for several more steps (Fig. 1c, right).

To determine whether the learned visuomotor mapping generalized to the left leg/foot, we compared the mean foot-placement error during the last ten baseline trials (when using the left foot to step to the target) to the foot-placement error during the generalization trial. We found that both the control and consequence groups generalized the learned mapping to the non-adapted left foot (mixed-model ANOVA, Group x Phase interaction: F_1,22_ = 12.70, p = 0.002, 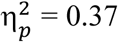), reflected by foot-placement errors in the direction opposite to the learned prism shift (i.e., a negative value) (Fig. 4a). However, the foot-placement error during the generalization trial differed significantly between groups, such that the consequence group demonstrated greater leftward deviation in foot placement from the target (see also Fig. S3), indicating greater generalization to the left leg/foot during precision walking.

**Figure 4.**
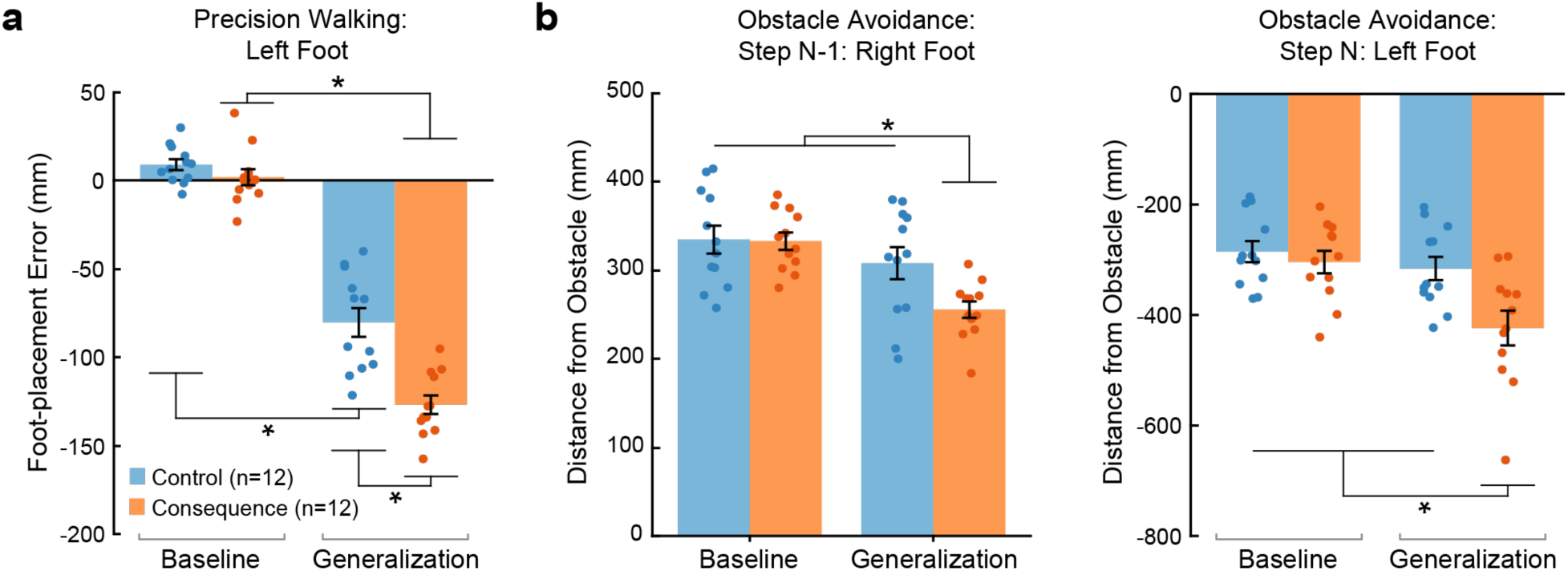
Generalization. ***a)*** Group mean ± SE foot-placement error for the baseline (mean of the last ten trials) and generalization phases during the precision walking task for the control (blue) and consequence (orange) groups. Foot-placement errors in the direction opposite to the prism shift (i.e., a negative value) indicate generalization. ***b)*** Group mean ± SE foot-placement error for the baseline (mean of the last ten trials) and generalization phases during the obstacle avoidance task for both the trailing foot (i.e., step N-1: right foot) and leading foot (i.e., step N: left foot) for the control (blue) and consequence (orange) groups. A smaller value indicates generalization for step N-1 (right foot), whereas a greater negative value reflects generalization for step N (left foot). Individual participant values are superimposed (n = 12 per group). * Indicate that values are significantly different from each other based on post hoc tests (p < 0.05). See Fig. S3 for more detailed post hoc test results.

For the obstacle avoidance task, we compared foot-placement deviation from the obstacle during the baseline phase (mean of last ten trials) and the generalization trial for both the trailing foot (i.e., step N-1: right foot) and leading foot (i.e., step N: left foot). For step N-1 (right foot), a smaller value reflects less deviation of the foot from the obstacle and indicates generalization of the learned mapping to the trailing right foot. We found that the consequence group demonstrated a smaller deviation of the right foot from the obstacle (i.e., a leftward shift in foot placement) during step N-1 compared to the control group (Fig. 4b and Fig. S3; mixed-model ANOVA, Group x Phase interaction: F_1,22_ = 10.98, p = 0.003, 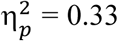). For step N (left foot), a greater negative value reflects greater deviation of the foot from the obstacle and indicates generalization of the learned mapping to the leading left foot. We found that the consequence group demonstrated greater deviation of the leading left foot from the obstacle (Fig. 4b and Fig. S3; mixed-model ANOVA, Group x Phase interaction: F_1,22_ = 16.18, p = 0.0006, 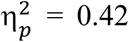). Taken together, experiencing an adverse physical consequence when making movement errors increases the degree of generalization across different visually guided walking tasks.

### Greater motor memory consolidation occurred in the group who experienced the adverse physical consequence

We also probed consolidation, defined here as memory stabilization of the learned prism-induced mapping such that it is resistant to retrograde interference by another (competing) mapping (Krakauer et al. 2005). Thus, at the end of the first testing session, participants performed 60 trials of the same adaptation task (i.e., precision walking with the right foot stepping on the target) while wearing the 20-diopter lenses that shifted the visual field in the opposite (i.e., leftward) direction of mapping A (which we refer to as mapping B; Fig. 1d). Following the initial testing session, the participants returned to the lab one week later to probe motor memory consolidation. Participants first performed 20 baseline trials of the adaptation task while wearing the zero-diopter lenses. Thereafter, all participants repeated the 60 adaptation trials with the 20-diopter (rightward-shifting) prism lenses to assess consolidation of mapping A. There was no slippery surface (and hence no adverse physical consequence for making a movement error) during the second testing session.

To determine the presence of consolidation, we compared three measures across testing sessions: the first adaptation trial error (representing the initial recall of the mapping), early adaptation error (i.e., mean of adaptation trials 2 – 8), and rate of adaptation. We used a single exponential for our adaptation rate measure. The R^2^ values of the individual fits in session 1 for the control group were 0.77 ± 0.09 (range: 0.60 to 0.90) and the consequence group were 0.79 ± 0.10 (range: 0.62 to 0.91). The R^2^ values of the individual fits in session 2 for the control group were 0.68 ± 0.16 (range: 0.38 to 0.89) and the consequence group were 0.53 ± 0.12 (range: 0.37 to 0.76). Figure 5a illustrates group mean foot-placement error during the baseline and adaptation phases during both testing sessions for the control and consequence groups. We found that the consequence group demonstrated greater error reduction in the first adaptation trial compared to the control group (mixed-model ANOVA, Group x Session interaction: F_1,22_ = 18.18, p = 0.0003, 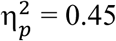), reflecting greater recall of the learned mapping one week later (Fig. 5b and Fig. S4).

**Figure 5.**
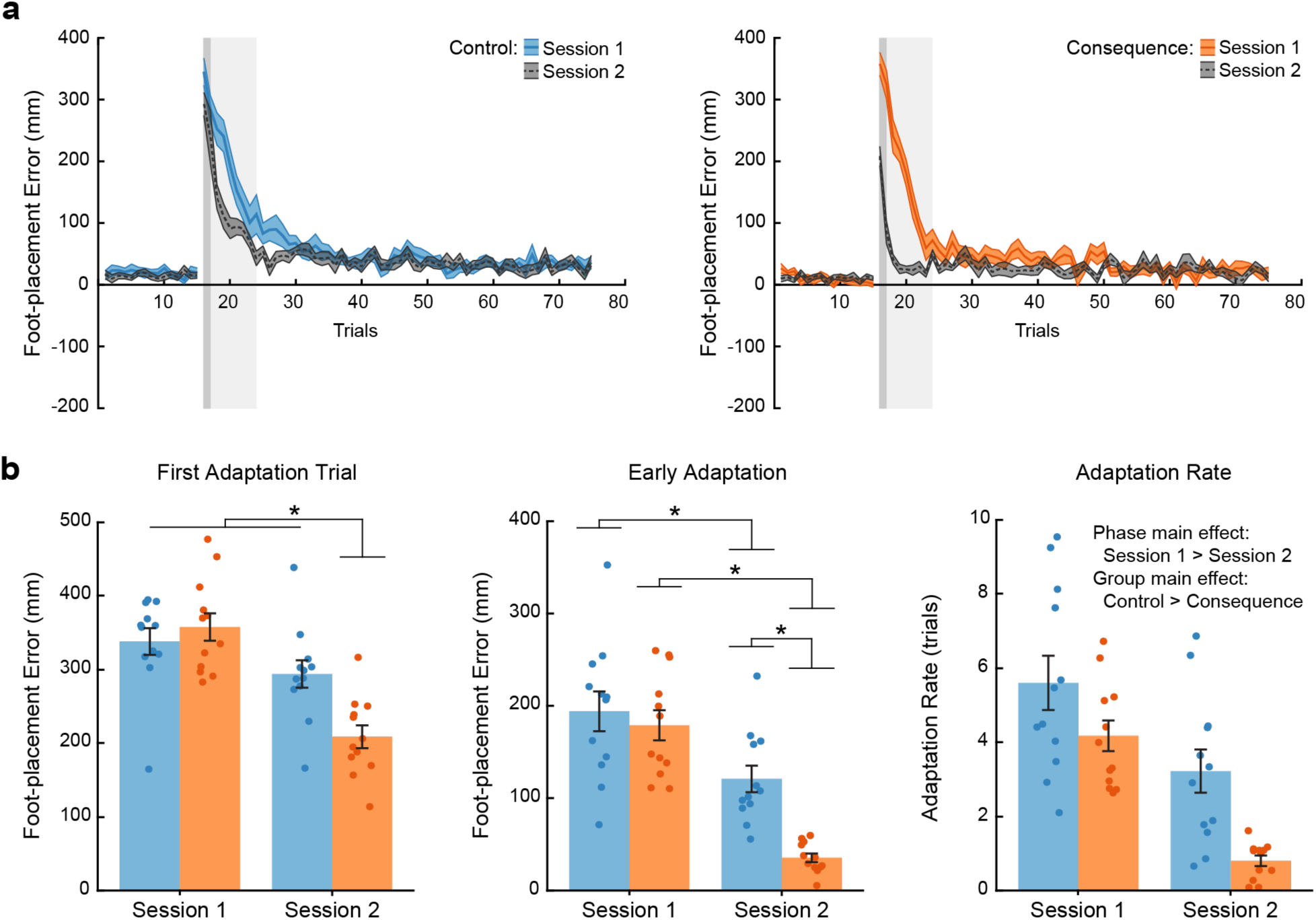
Motor memory consolidation. ***a)*** Group mean ± SE foot-placement error for all trials in the baseline and adaptation phases across testing sessions for the control (blue) and consequence (orange) groups (n = 12 per group for both sessions). ***b)*** Group mean ± SE for the first adaptation trial error (dark grey shaded box in panel a), early adaptation error (light grey shaded box in panel a), and rate of adaptation across testing sessions for the control (blue) and consequence (orange) groups. One week separated testing sessions. Individual participant values are superimposed (n = 12 per group). * Indicate that values are significantly different from each other based on post hoc tests (p < 0.05). See also Fig. S4 for more detailed post hoc test results.

A faster reduction in foot-placement error (i.e., savings) indicates that the learned mapping was consolidated. To quantify savings of the learned mapping, we compared foot-placement error during early adaptation (i.e., mean of adaptation trials 2 – 8) and the rate of adaptation across testing sessions. We found that both the control and consequence groups showed a significant reduction in foot-placement error during early adaptation (Fig. 5b and Fig. S4; mixed-model ANOVA, Group x Session interaction: F_1,22_ = 7.98, p = 0.010, 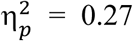). However, the consequence group demonstrated significantly greater error reduction during the second testing session compared to the control group, reflecting greater savings one week later. We also found that both the control and consequence groups demonstrated a faster rate of relearning during the second testing session (Fig. 5b and Fig. S4; mixed-model ANOVA, Session main effect: F_1,44_ = 30.99, p = 1.454e-6, 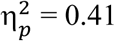). In addition, we found that the consequence group demonstrated a faster rate of adaptation across both testing sessions (mixed-model ANOVA, Group main effect: F_1,44_ = 13.86, p = 0.0006, 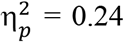), though we did not detect a significant Group x Session interaction for this measure (mixed-model ANOVA, F_1,44_ = 0.92, p = 0.344, 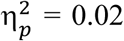). Takentogether, experiencing an adverse physical consequence when making movement errors increases initial recall and savings of a learned mapping one week later, reflective of greater motor memory consolidation.

## DISCUSSION

Learning from the consequences of our actions is imperative for safe and successful motor performance. To determine which behaviors to maintain, people presumably learn to dissociate actions that give rise to desirable outcomes from those that do not. Here we tested the hypothesis that experiencing a balance-threatening physical consequence when making a movement error serves to enhance motor memory. Previous research on reaching adaptation has reported limited generalization (Ghahramani et al. 1996; Krakauer et al. 2000; Wang 2008) and mixed evidence for motor memory consolidation (Brashers-Krug et al. 1996; Caithness et al. 2004; Krakauer et al. 2005)—though research on walking has found stronger evidence of both (Alexander et al. 2011, 2013; Maeda et al. 2017b, 2018; Malone et al. 2011). In this study, we found that participants who experienced an adverse physical consequence when making foot-placement errors during adaptation demonstrated increased interlimb-transfer on a precision walking task and generalization to an obstacle-avoidance task. Furthermore, this group showed increased recall and savings of the learned visuomotor mapping one week later despite exposure to a competing mapping during initial learning—evidence of greater motor memory consolidation. The differences in generalization and consolidation between groups occurred even though they both experienced similar magnitude foot-placement errors and adapted at similar rates. Our results suggest that the brain considers the potential physical consequences of movement error when learning and that balance-threatening consequences serve to enhance this process.

Reward and punishment are known to reinforce motor learning. If one considers our physical consequence as punishment, then our results contrast with previous work that shows punishment accelerates adaptation rate but has little effect on later retention (Galea et al. 2015; Song and Smiley-Oyen 2017; Song et al. 2020). On the other hand, avoiding the adverse physical consequence may serve as a reward. Specifically, as participants adapt to the prisms and become more accurate stepping to the target, they decrease the likelihood of contacting the slippery surface and suffering a potential loss of balance. Thus, foot placement on the target, or foot placement with minimal error such that the foot does not hit the slippery surface, may act as reward-like feedback. Interestingly, avoiding an aversive outcome causes activation in a brain region—medial orbitofrontal cortex—also implicated in encoding reward (Kim et al. 2006). In this case, our results are compatible with previous work that shows reward enhances retention but does not affect adaptation rate (Galea et al. 2015; Song and Smiley-Oyen 2017), though it is important to note that reward can accelerate learning depending on the reward structure (Nikooyan and Ahmed 2015). However, it may not be appropriate to associate our physical consequence with monetary reward or punishment per se. Rather, the slip serves as a functionally meaningful consequence of movement error, thus making it a more ecological manipulation.

Perhaps surprisingly, we found similar adaptation rates between groups in session 1 even though the one group experienced the adverse physical consequence. However, there is minimal room for differences given that rates are already quite fast in this paradigm (i.e., less than six trials on average). Thus, the rapid nature of adaptation may have masked any potential group differences. In addition, the slip exposure does not provide information on how to adapt to the new prism-induced visuomotor mapping, which may further explain the lack of differences. Alternatively, our results suggest that the adverse physical consequence may simply influence the strength of the learned mapping. Specifically, since the control of balance is fundamental to movement, the brain may assign greater importance (or value) to maintaining the learned visuomotor mapping because it ensures the slip is avoided. This notion lends support to our previous work where we proposed that the threat of falling during balance-challenged adaptation tasks may have contributed to our findings of greater motor learning (Bakkum et al. 2020, 2021). Thus, just like expected value increases with the probability of reward, value may increase with the probability of maintaining balance.

Our physical consequence threatened balance, and the unexpected nature of at least the first slip experience likely surprised participants. Both threat and surprise can increase emotional arousal. We propose that experiencing the adverse physical consequence when making foot-placement errors may have enhanced motor memory through increased recruitment of brain regions engaged in processing emotional arousal. Research in humans and other animals provides compelling evidence for the role of the amygdala in forming and maintaining lasting memories associated with emotional arousal (McGaugh 2004). For example, in humans, the threat of being shocked enhances declarative memory through activation of the amygdala (Murty et al. 2012). Furthermore, lesions to the amygdala attenuate the advantageous effects of emotional arousal on memory (McGaugh et al. 1996). The locus coeruleus (LC), which is heavily connected to the amygdala, is also activated in response to emotionally arousing stimuli, including threat, and can facilitate memory encoding (Clewett et al. 2018; Tully and Bolshakov 2010). In addition, the anterior cingulate cortex (ACC) is active in response to error detection or surprise (Hayden et al. 2011) and interestingly, shows greater electroencephalography-based theta band spectral power following a loss of balance during walking (Sipp et al. 2013). The amygdala, LC, and ACC connect directly or indirectly to the motor cortex, cerebellum, and basal ganglia (Farley et al. 2016; Grèzes et al. 2014; Rolls 2019; Schönfeld and Wojtecki 2019; Tully and Bolshakov 2010), which are each implicated in motor memory consolidation (Debas et al. 2010; Landi et al. 2011; Leow et al. 2012). Thus, the emotionally charged experience of slipping may have increased the activation of one or more of these regions during adaptation and led to strengthening of synaptic connections in relevant sensorimotor areas where memory of the learned mapping was marked for consolidation. This idea resembles the emotional tagging hypothesis (McReynolds and McIntyre 2012; Richter-Levin and Akirav 2003), which attempts to explain how and why emotionally arousing events are better remembered.

The effects of the adverse physical consequence associated with movement error may derive from an implicit and/or explicit learning process. There is strong evidence that implicit, internal-model-based learning occurs in our walking adaptation paradigm (Maeda et al. 2017a). However, recent work highlights the contribution of explicit strategies to reinforcement-based visuomotor adaptation (Codol et al. 2018; Holland et al. 2018; Song et al. 2020). Given the nature of our physical consequence, it may draw greater attention to the error and thus, serve to increase the reliance on using an explicit aiming strategy to regain movement accuracy. We suggest that it is likely a combination of both implicit- and explicit-based learning, but the distinction between these processes is beyond the scope of this study.

Overall, our findings show that experiencing an adverse physical consequence when making errors, which threatens stability, enhances sensorimotor learning. Specifically, the consequence group better generalized learning to different walking tasks and showed greater consolidation one week later. This may suggest our motor systems are tuned to remember behaviors that promote personal safety and provide an important survival advantage (Nairne et al. 2007). Our work highlights a new factor that affects sensorimotor learning and provides an interesting avenue for future research. Our findings also provide intriguing implications for neurorehabilitation aimed at long-lasting performance improvements that generalize beyond a clinical setting. They suggest that therapists should consider incorporating tasks or situations that elicit a threatening physical consequence if the patient moves incorrectly or in manner inconsistent with how a therapist is training them. Safety is paramount in these cases, which can be managed using fall safety harness systems and/or virtual reality.

## AUTHOR CONTRIBUTIONS

A.B. and D.S.M. conceived and designed research; A.B. performed experiments; A.B. analyzed data; A.B. and D.S.M. interpreted results of experiments; A.B. prepared figures; A.B. and D.S.M. drafted manuscript; A.B. and D.S.M. edited and revised manuscript; A.B. and D.S.M. approved final version of manuscript.

