## Supplemental Figures for "Learning from the physical consequences of our actions improves motor memory"

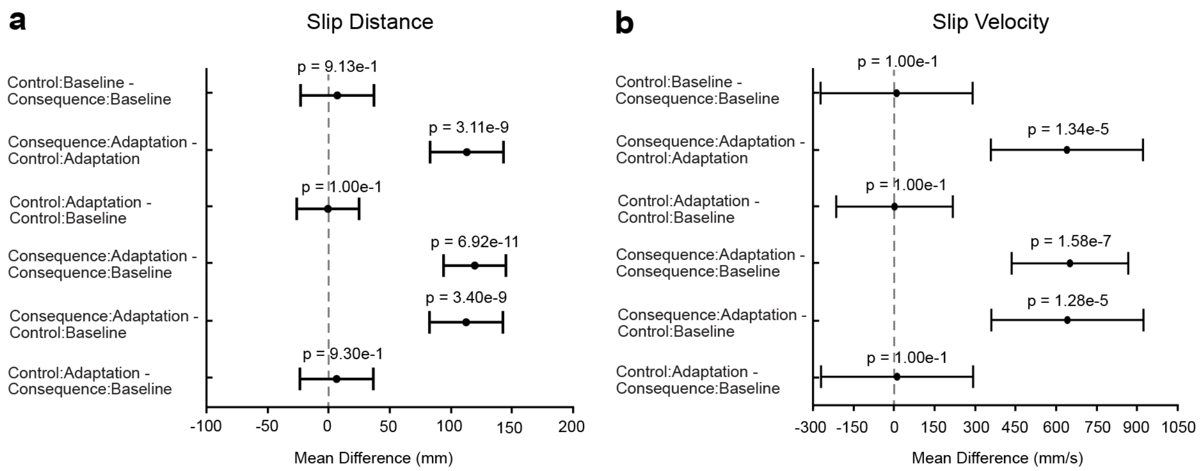


**Figure S1. Least square means Tukey honestly significant difference tests of pairwise comparisons related to Figure 2.** (a) Pairwise comparisons for slip distance following a statistically significant Group x Phase interaction. (b) Pairwise comparisons for slip velocity following a statistically significant Group x Phase interaction. Values are shown as mean differences with 95% confidence intervals and p values.

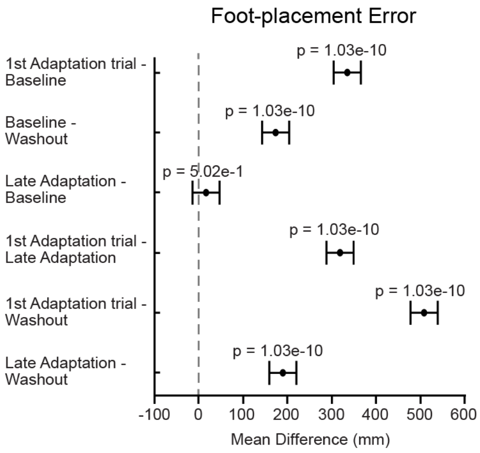


**Figure S2. Least square means Tukey honestly significant difference tests of pairwise comparisons related to Figure 3.** Pairwise comparisons for foot-placement error following a statistically significant main effect of Phase. Values are shown as mean differences with 95% confidence intervals and p values.

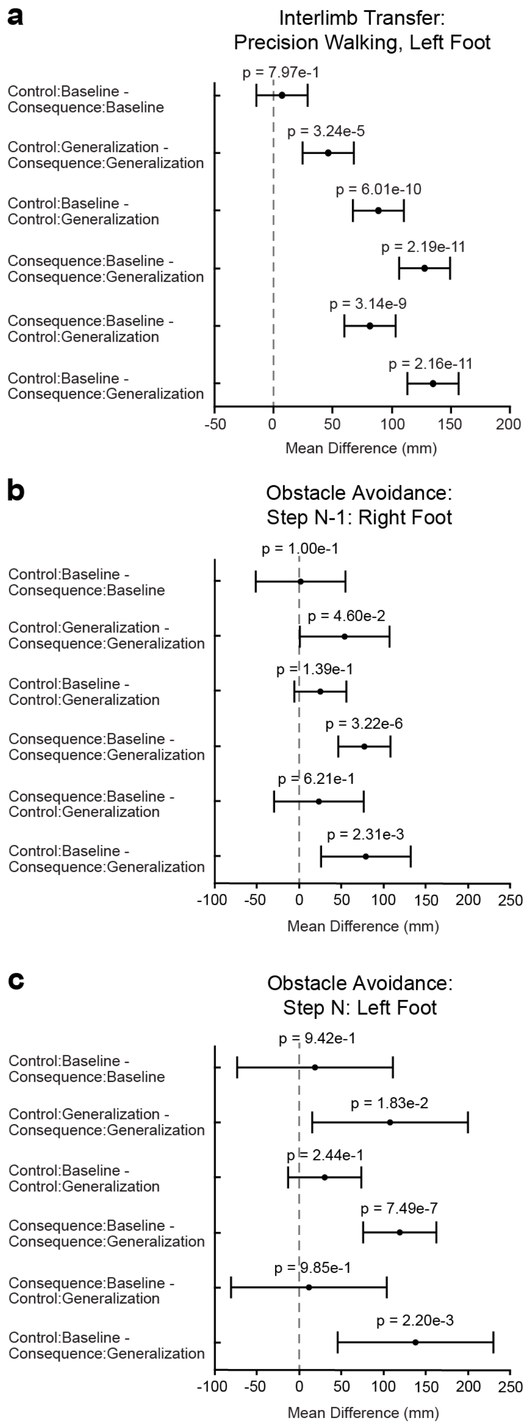


**Figure S3. Least square means Tukey honestly significant difference tests of pairwise comparisons related to Figure 4.** (a) Pairwise comparisons for foot-placement error in the interlimb transfer test following a statistically significant Group x Phase interaction. (b) Pairwise comparisons for foot placement of step N-1 (right foot) relative to the obstacle following a statistically significant Group x Phase interaction. (c) Pairwise comparisons for foot placement of step N (left foot) relative to the obstacle following a statistically significant Group x Phase interaction. Values are shown as mean differences with 95% confidence intervals and p values.

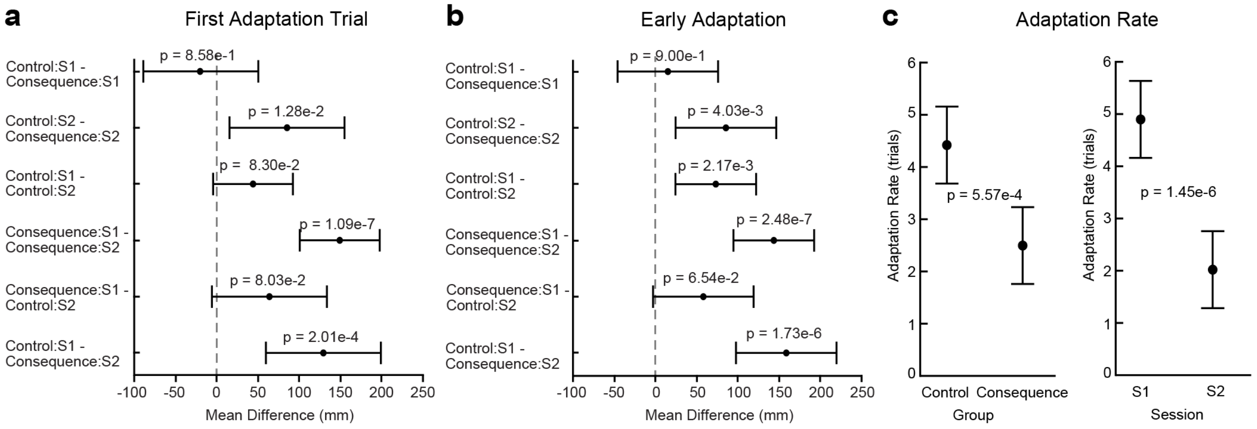


**Figure S4. Least square means Tukey honestly significant difference tests of pairwise comparisons and least square means plots related to Figure 5.** (a) Pairwise comparisons for first adaptation trial foot-placement error following a statistically significant Group x Session interaction. (b) Pairwise comparisons for early adaptation foot-placement error following a statistically significant Group x Session interaction. (c) Least square means plots for adaptation rate following statistically significant main effects. Values are shown as mean differences with 95% confidence intervals and p values for (a) and (b). Values are shown as least square means with 95% confidence intervals and p values for (c). S1 = session 1; S2 = session 2.
